# Co-entangled actin-microtubule composites exhibit tunable stiffening and power-law stress relaxation

**DOI:** 10.1101/262089

**Authors:** Shea N. Ricketts, Jennifer L. Ross, Rae M. Robertson-Anderson

## Abstract

We use optical tweezers microrheology and fluorescence microscopy to characterize the nonlinear mesoscale mechanics and mobility of in vitro co-entangled actin-microtubule composites. We create a suite of randomly-oriented, well-mixed networks of actin and microtubules by co-polymerizing varying ratios of actin and tubulin in situ. To perturb each composite far from equilibrium, we use optical tweezers to displace an embedded microsphere a distance greater than the lengths of the filaments at a speed much faster than their intrinsic relaxation rates. We simultaneously measure the resistive force the filaments exert and the subsequent force relaxation. We find that the presence of a large fraction of microtubules (>0.7) is needed to substantially increase the resistive force, which is accompanied by large heterogeneities in force response. Actin minimizes these heterogeneities by reducing the mesh size of the composites and supporting microtubules against buckling. Composites also undergo a sharp transition from stress-softening to stiffening when the fraction of microtubules (*ϕ*_*T*_) exceeds 0.5, by microtubules suppressing actin bending fluctuations. The induced force following strain relaxes via two time-dependent power-law decays. The first decay phase, with scaling exponents that increase proportionally with the fraction of actin, signifies actin bending fluctuations. Alternatively, the second phase, with a *ϕ*_*T*_-independent scaling exponent of ~0.4, is indicative of filaments reptating out of deformed entanglement constraints. Corresponding mobility measurements of steady-state actin and microtubules show that both filaments are more mobile in equimolar composites (*ϕ*_*T*_=0.5) compared to networks of primarily actin or microtubules. This non-monotonic dependence of mobility on *ϕ*_*T*_, which further demonstrates the important role mesh size plays in composites, highlights the surprising emergent properties that can arise in composites.

## Introduction

The cytoskeleton is a complex network of protein filaments that gives eukaryotic cells structural integrity and shape while enabling cell motility, division and morphogenesis. Such multifunctional mechanics and processes are possible because of the varying structural properties of cytoskeletal proteins, as well as the interactions between them (1–4). Actin and tubulin are two such proteins that form filaments with very different stiffnesses. Tubulin dimers (~110 kDa) polymerize into 25 nm wide hollow, rigid microtubules with persistence lengths of *l*_*p*_ ≈ 1 mm, whereas actin monomers (~42 kDa) polymerize into ~7 nm wide semiflexible filaments (F-actin) with *l*_*p*_ ≈ 10 µm (5, 6). At high concentrations, both actin filaments and microtubules form sterically interacting entangled networks with filament dynamics that can be described by the reptation or tube model. Within this framework, each filament is confined to a tube-like region formed by surrounding constraining filaments, and is forced to relax induced stress by diffusing curvilinearly (i.e. reptating) out of its deformed tube (7, 8). The timescale for this process, often termed disengagement, is on the order of minutes to hours for actin and microtubules (9–11). Entangled actin can also partially relax via faster mechanisms such as bending fluctuations (12–14). The degree of filament entanglement and density in networks of actin and microtubules can be characterized by their respective mesh sizes, 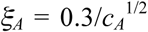 and 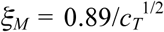, where *c*_*A*_ and *c*_*T*_ are the corresponding protein concentrations in units of mg/ml and the resulting mesh sizes are in units of microns (15–17). As can be seen from these equations, because of the differences in molar mass and filament structure of the two proteins, the mesh size for a microtubule network of a given molarity is ~2*x* larger than that for actin.

Steric interactions between actin and microtubules within cells directly influence cell shape and mechanics by regulating filament mobility and providing coordinated reinforcement to the cytoskeleton (6, 18). Processes such as cell motility and cytokinesis also rely on the physical interactions between these filaments (6, 19). Further, while individual microtubules buckle under substantial compressive forces (20–23), steric interactions with the elastic actin network allow microtubules to bear larger compressive loads within the cell (6, 10, 20, 24). Studies of composite networks of actin and microtubules are further motivated by materials engineering, in which soft elastic networks are often reinforced with stiff fibers or rigid rods. By tuning the concentration of rigid rods relative to the flexible filaments, bulk properties of composites can be optimized to create lightweight materials with high strength and stiffness (19, 25). Such composites also offer enhanced control over large-scale mechanics and increased failure limits, tuned by the mechanical differences of the two composite constituents. Finally, composites both in nature and in industry often exhibit emergent properties in which the resulting mechanical properties are not a simple sum of the single-component network mechanics.

In vitro studies of cytoskeleton networks have largely focused on single-component systems of either actin or microtubules (15, 26–28). One previous passive microrheology study of an equimolar composite of actin and microtubules showed that while entangled actin solutions appeared incompressible, the composite displayed a finite compressibility resulting from the high stiffness of microtubules (10, 29). This same study showed that the crossover to viscous behavior that microtubule networks exhibit at low strain rates was not observed for actin solutions or equimolar composites. A previous nonlinear bulk rheology study of crosslinked actin networks doped with low concentrations of microtubules (~3 – 10*x* lower than the actin concentration) showed that the addition of microtubules led to nonlinear strain-stiffening as compared to the signature strain-softening behavior of entangled and weakly-crosslinked actin networks (19, 30, 31). The authors explained this shift as due to stiff microtubules suppressing actin bending modes and local fluctuations, leading to enhanced stretching and affine deformation. While these few studies revealed important emergent properties in actin-microtubule composites, they were limited in the parameter space of composite makeup. Thus, the question remains as to how the relative concentrations of actin and microtubules impact the mechanical properties of composites. Further, these studies probed the bulk response resulting from large-scale nonlinear strains, and the microscopic linear response due to passively diffusing microspheres. Yet, the relevant lengthscales in actin-microtubule composites are in between these two scales, with persistence lengths of ~10 µm to 1 mm, filament lengths of ~5 – 20 µm, and mesh sizes on the order of a micron. Finally, in these prior studies, composites were created by adding pre-polymerized microtubules to actin, rather than polymerizing both proteins together. This method often induces flow alignment of filaments, shearing of microtubules, and bundling of actin filaments, preventing the formation of truly isotropic, well integrated, co-entangled composites.

Here, we create co-polymerized, co-entangled actin-microtubule composites with systematically varying actin:tubulin molar ratios. We characterize the nonlinear mesoscale mechanics of composites by pulling optically-trapped microspheres a distance of 30 µm through the composites at a rate much faster than the system relaxation rates while measuring the local force induced in the composites. These measurements perturb the composites far from equilibrium, and are uniquely able to probe possible buckling, rupture, and rearrangement events, as well as micro- and mesoscale spatial heterogeneities in the composites. We complement these nonlinear measurements with image analysis of steady-state composites.

We find that composites with more microtubules than actin (*ϕ*_*T*_>0.5) exhibit stress-stiffening as opposed to the softening displayed in actin-dominated composites. The onset of stiffening is coupled with a large drop in steady-state filament mobility. We also find that high concentrations of microtubules (*ϕ*_*T*_>0.7) are needed to substantially increase resistive force and induce more pronounced heterogeneities in force response, which we show arises from the smaller mesh size of actin relative to microtubules. This mesh size mismatch also leads to an emergent non-monotonic dependence of filament mobility on *ϕ*_*T*_, with equimolar composites (*ϕ*_*T*_=0.5) displaying the most pronounced steady-state mobility. Following strain, the induced force relaxes via two power-law decays for all composites. The initial fast decay, arising from actin bending modes, is increasingly suppressed as the relative concentration of microtubules increases. Conversely, the second phase slow decay is robust to changes in network composition and is indicative of entangled filaments diffusing out of non-classical entanglement tubes.

## Materials and Methods

Rabbit skeletal actin, porcine brain tubulin, and rhodamine-labeled tubulin were purchased from Cytoskeleton (AKL99, T240, TL590M), and Alexa-488-labeled actin was purchased from Thermofisher (A12373). To form actin-microtubule composites, varying ratios of unlabeled actin monomers and tubulin dimers were suspended in an aqueous buffer containing 100 mM PIPES, 2 mM MgCl_2_, 2 mM EGTA, 1 mM ATP, 1 mM GTP, and 5 µM Taxol to a final total protein concentration of 11.3 µM (Fig 1A). This buffer was optimized to achieve co-polymerization and stabilization of actin and microtubules. To image composite architecture, 0.13 µM of pre-assembled Alexa-488-labeled actin filaments, at a 1:1 labeled:unlabeled monomer ratio, and 0.19 µM pre-assembled rhodamine-labeled microtubules, at a 1:5 labeling ratio, were also added to the solution as tracer filaments (Fig 1B). For microrheology measurements (Fig 1C,D), a sparse amount of 4.5 µm microspheres (probes, Polysciences, Inc.), coated with Alexa-488 BSA (Invitrogen), were added. This protein and bead mixture was pipetted into a sample chamber made from a glass slide and coverslip, separated by ~100 µm with double-sided tape to accommodate ~20 µL, and sealed with epoxy. Co-polymerization of both proteins was achieved by incubating the sample for 1 hour at 37 ºC prior to measurement, resulting in a well-integrated and stable co-entangled composite (Fig S1 in the Supplemental Materials). The molar fraction of tubulin in composites, *ϕ*_*T*_ = [*tubulin*]/([*actin*]+[*tubulin*]) where [*actin*] and [*tubulin*] are the molar concentrations of each protein, was varied from 0 to 1 while total protein concentration (labeled and unlabeled actin and microtubules) was held fixed at 11.6 µM.

**Figure 1.**
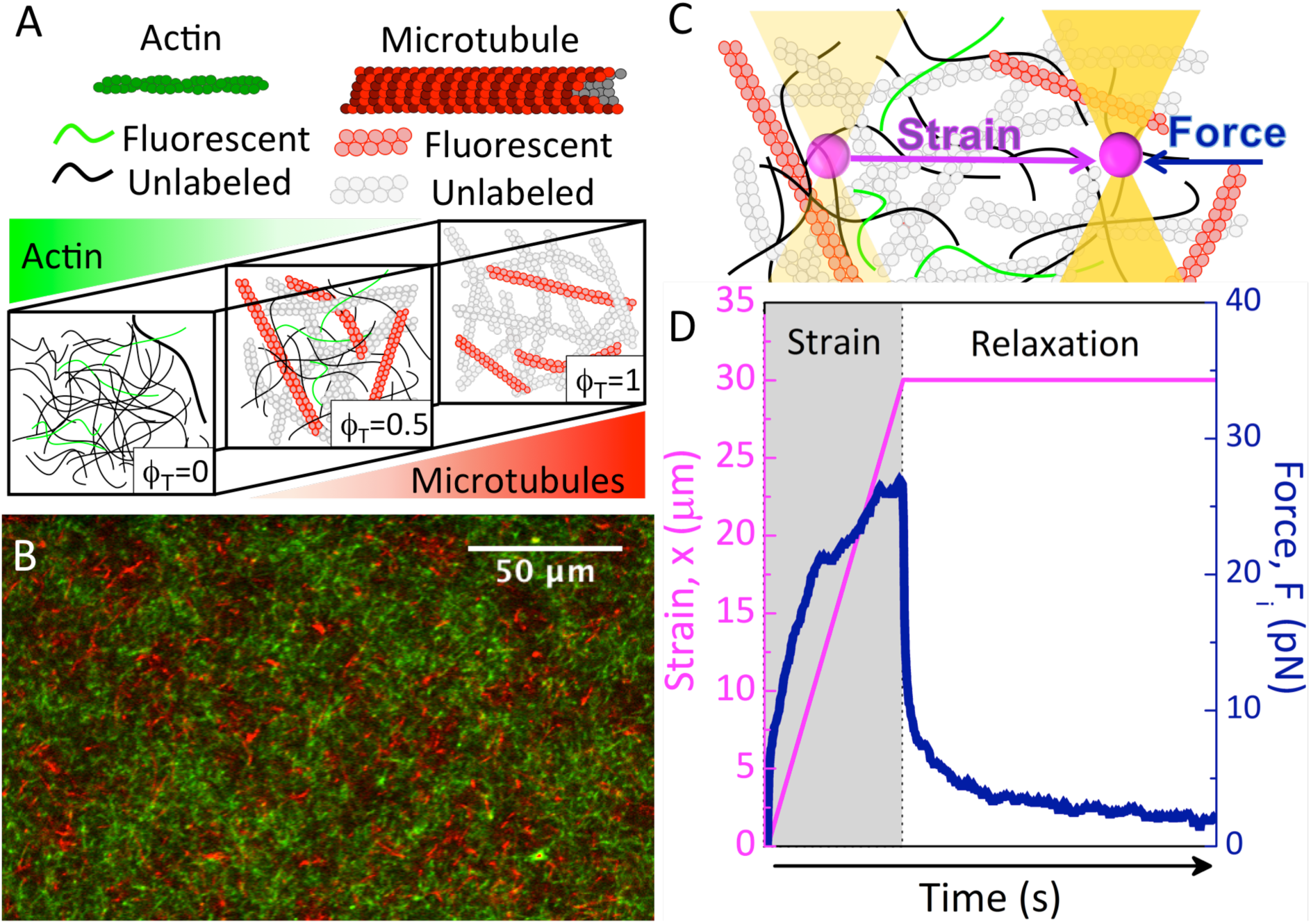
Schematic of experimental approach. (A) Cartoon of molecular components used in actin-microtubule composites. The tubulin molar fraction, *ϕ*_*T*_ = [*tubulin*]/([*actin*]+[*tubulin*]), is varied from 0 to 1 with total protein concentration fixed at 11.6 µM. (B) To display network architecture, 0.13 µM of actin and microtubules are labeled with Alexa-488 (green) and rhodamine (red), respectively. The image shown is a two-color laser scanning confocal image of a *ϕ*_*T*_ = 0.5 composite. (C) For microrheology measurements an optically-trapped microsphere probe embedded in the composite is displaced a distance *x* = 30 µm (Strain, magenta) at a speed of 20 µm/s while the resistive force, *F_i_(x,t)*, that the composite exerts on the probe is measured (Force, blue). (D) Sample microrheology data showing the resistive force (blue) and probe position (magenta) during (Strain) and following (Relaxation) probe displacement. Data shown is for the *ϕ*_*T*_ = 0.5 composite.

The optical trap used in microrheology measurements was formed using an IX71 fluorescence microscope (Olympus) outfitted with a 1064 nm Nd:YAG fiber laser (Manlight) focused with a 60*x* 1.4 NA objective (Olympus). A position-sensing detector (Pacific Silicon Sensor) measured the deflection of the trapping laser, which is proportional to the force acting on the trapped probe. Trap stiffness was calibrated using the Stokes drag method (32–34). During force measurements, a probe embedded in the composite was trapped and moved a distance *x* = 30 µm at a constant speed of 20 µm/s relative to the sample chamber using a nanopositioning piezoelectric stage (Mad City Labs) (Fig 1C,D). Laser deflection and stage position were recorded at a rate of 20 kHz using custom-written Labview code. Post-measurement data analysis was done using custom-written MATLAB software. Displayed force curves in Figures 2 and 4 are averages over an ensemble of 25 different trials, *i*, with each *i*^th^ trial located in a different region of the network separated by >100 µm.

**Figure 2.**
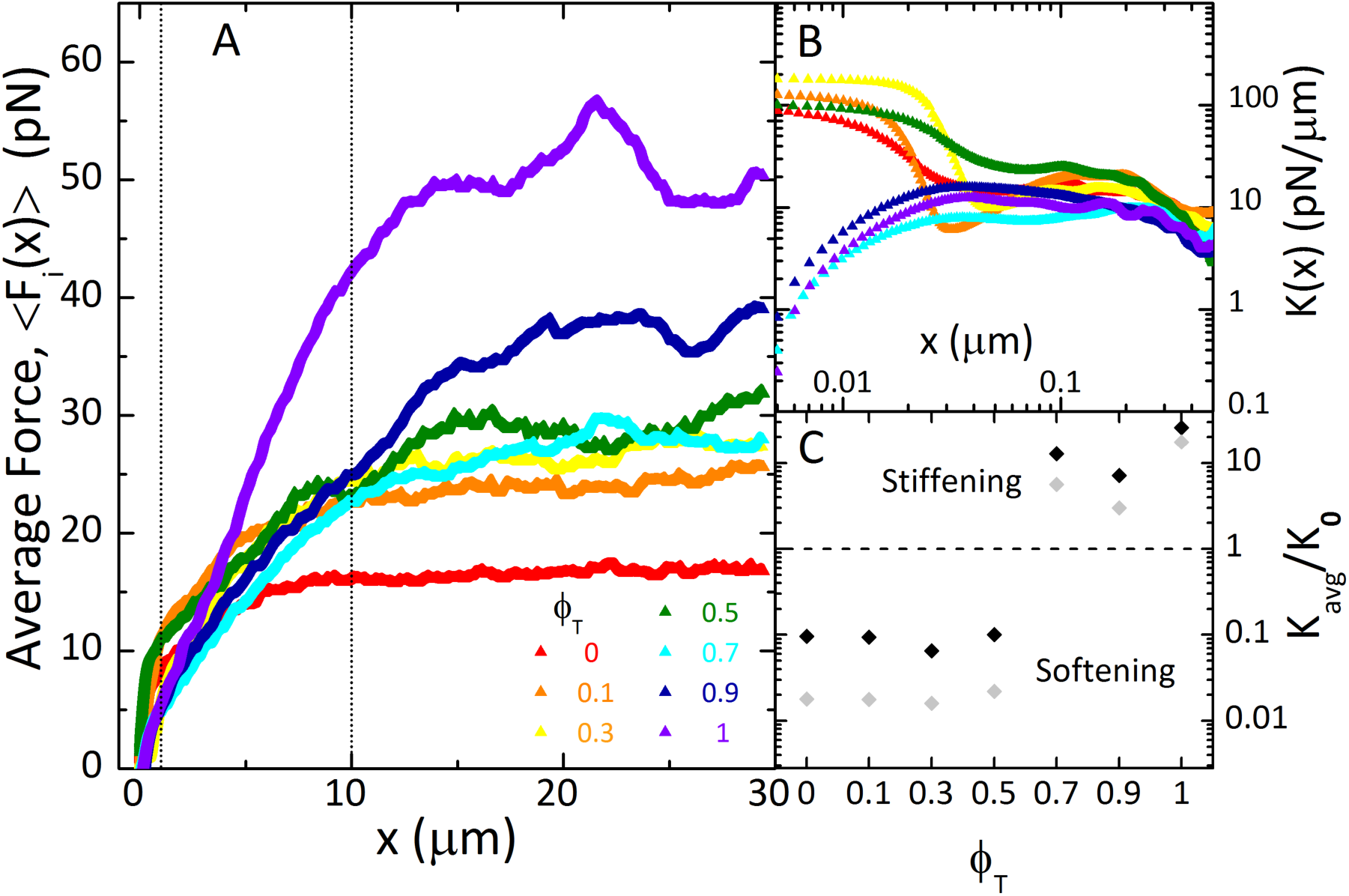
Mesoscale force response of actin-microtubule composites display a *ϕ*_*T*_-dependent crossover from stress-softening to stress-stiffening. (A) Ensemble-averaged resistive force, *<F_i_(x)>*, exerted on moving probe by actin-microtubule composites of varying tubulin molar fraction *ϕ*_*T*_ (listed in legend). Data shown is an average over 25 individual, *i*, measurements. (B) Differential modulus, *K(x)*, calculated as the derivative of the average force with respect to strain distance *x* (*K(x) = d<F_i_(x)>/dx*). (C) Differential modulus values averaged over strain distances of *x* = 1 µm (black) and *x* = 10 µm (grey), and normalized by the corresponding value at *x* = 0 (*K*_*0*_) for each composite (*K*_*avg*_/*K*_*0*_). Strain distances averaged over are shown as dotted vertical lines in A. As shown, composites transition from stress-softening (*K*_*avg*_/*K*_*0*_ < 1) to stress-stiffening (*K*_*avg*_/*K*_*0*_ > 1) when the tubulin molar fraction *ϕ*_*T*_ exceeds 0.5.

To visualize network mobility (Fig 5) and architecture (Fig S1), we used a Nikon A1R laser scanning confocal microscope with 60*x* objective to collect 2D images, 2D time-series, and 3D images of composites. The microscope has 488 nm and 532 nm lasers to alternately record separate images in green and red channels to visualize 488-actin and rhodamine-tubulin, respectively. To characterize filament mobility within composites, 512×512 pixel time-series with pixel size of 0.41 µm were recorded at 30 fps for 3 minutes in green and red channels. Post-imaging analysis was carried out using ImageJ/Fiji. Color channels were separated upon import and analyzed separately. To reduce noise, each time-series was averaged over every 30 frames, resulting in 1 fps time-series of 180 frames. We then computed the standard deviation (SD) of intensity values over time for each pixel of the time-series, and displayed the result as a single 512x512, 32-bit image which we call the “mobility image” (Fig 5A). In these images, SD intensity correlates with filament mobility such that a low (black) SD results from a filament remaining in the same position over time (no change in pixel intensity over time) while a high (white) SD indicates high filament mobility (pixel intensity changing a lot over time). The intensity of each pixel of the mobility image was then averaged over the entire image (44 µm^2^) to give a spatial average of the standard deviation for each time-series (<SD>). We performed this measurement on 3–5 movies for each network composition (*ϕ*_*T*_). The average standard deviation <SD> and the corresponding standard error was computed for each *ϕ*_*T*_ and plotted for the actin (green) and the microtubule (red) channels separately (Fig 5B).

## Results and Discussion

While actin and microtubules coexist in cells, the standard in vitro polymerization and network formation conditions for the two proteins are incompatible. The few previous studies that have investigated actin-microtubule networks have pre-polymerized and stabilized microtubules before adding to actin monomers to polymerize (3, 10), which can lead to flow alignment and rupturing of microtubules upon pipetting into actin monomer solutions and/or the experimental sample chamber. We instead sought to assemble unperturbed co-polymerized networks of actin and microtubules entangled with one another within an experimental sample chamber. To do so, we designed hybrid buffers and polymerization methods detailed in the Materials and Methods. Our methods result in isotropic, well-integrated composite networks of co-entangled actin and microtubules (Fig 1B, Fig S1).

We use active microrheology to measure the local force response of actin-microtubule composites with varying tubulin molar fractions, *ϕ*_*T*_, subject to nonlinear strains. The data presented in Figure 2 shows the ensemble-averaged force, *<F_i_(x)>*, exerted on optically trapped microspheres as they are pulled a distance *x* through each composite. As shown, introducing higher fractions of tubulin to composites increases resistive force (Fig 2A) and stiffness (Fig 2B,C). The differential modulus, *K(x) = d<F_i_(x)>/dx*, quantifies the degree of network stiffness, so increasing and decreasing *K(x)* values during strain signify stress stiffening and softening, respectively. As shown in Figure 2B, composites comprised of mostly actin (tubulin fraction *ϕ*_*T*_ ≤ 0.5) are initially relatively stiff but quickly soften. However, beyond *ϕ*_*T*_ = 0.5, composites undergo a stark transition, displaying an initially soft/viscous force response followed quickly by stress stiffening (Fig 2B). We can quantify the degree of stiffening/softening during the strain by averaging *K(x)* over different strain distances *x*, which we denote as *K*_*avg*_, and normalizing *K*_*avg*_ by the value measured at *x* = 0, *K*_*0*_ (Fig 2C). As shown, for strains up to *x* = 1 μm, the average differential modulus increases by an order of magnitude from its initial value for composites with mostly microtubules (*ϕ*_*T*_ ≥ 0.7), while it drops by an order of magnitude for *ϕ*_*T*_ ≤ 0.5 composites. As the strain continues (i.e. *x* increases) all composites exhibit increased softening as the force response approaches a viscous steady-state regime, as indicated by the measured forces approaching *x*-independent plateaus (Fig 2A) and the reduced *K*_*avg*_/*K*_*0*_ values for *x* = 10 μm compared to those for *x* = 1 μm (Fig 2C).

These results are qualitatively in line with previous bulk rheology results for crosslinked actin networks doped with microtubules. As described in the Introduction, these studies found that the addition of microtubules led actin networks to stress-stiffen rather than soften, due to microtubules suppressing actin bending modes (15, 19, 35). However, even for weakly-crosslinked actin networks, these studies found that microtubule fractions as low as *ϕ_T_ ≈* 0.3 induced stiffening (19), as opposed to *ϕ_T_ >* 0.5 in our experiments. Because crosslinking in actin networks can suppress bending modes and lead to stress stiffening (36), this difference likely arises from the lack of crosslinking in our composites. Further, it has been suggested that microtubules are able to propagate mechanical loads long distances across the cytoskeleton, enhancing the large-scale elastic response of otherwise flexible networks (29). Thus, the differences between our results and those of macrorheology may suggest that actin-microtubule composites have the ability to exhibit lengthscale-dependent nonlinear stress response, such as softening at the micro- and mesoscales but stiffening on a macroscopic scale. In fact, previous studies have shown that entangled actin networks can display scale-dependent nonlinear elasticity, exhibiting stress-stiffening at the microscale and stress-softening in bulk (37).

Increasing the fraction of tubulin in composites also increases the heterogeneities in force response (Fig 3). Microscale heterogeneities are displayed by increased noise or “bumpiness” in the individual force curves (Fig 3A), while large-scale heterogeneities can be seen by comparing the force curves for all *i* trials measured in different regions of the sample (separated by >100 um). Figure 3A, which shows the individual force curves, *F_i_(x)*, for all *i* measurements for actin networks (*ϕ*_*T*_ = 0), microtubule networks (*ϕ*_*T*_ = 1), and *ϕ*_*T*_ = 0.5 composites, clearly shows that increasing the fraction of microtubules increases heterogeneity at both scales.

**Figure 3.**
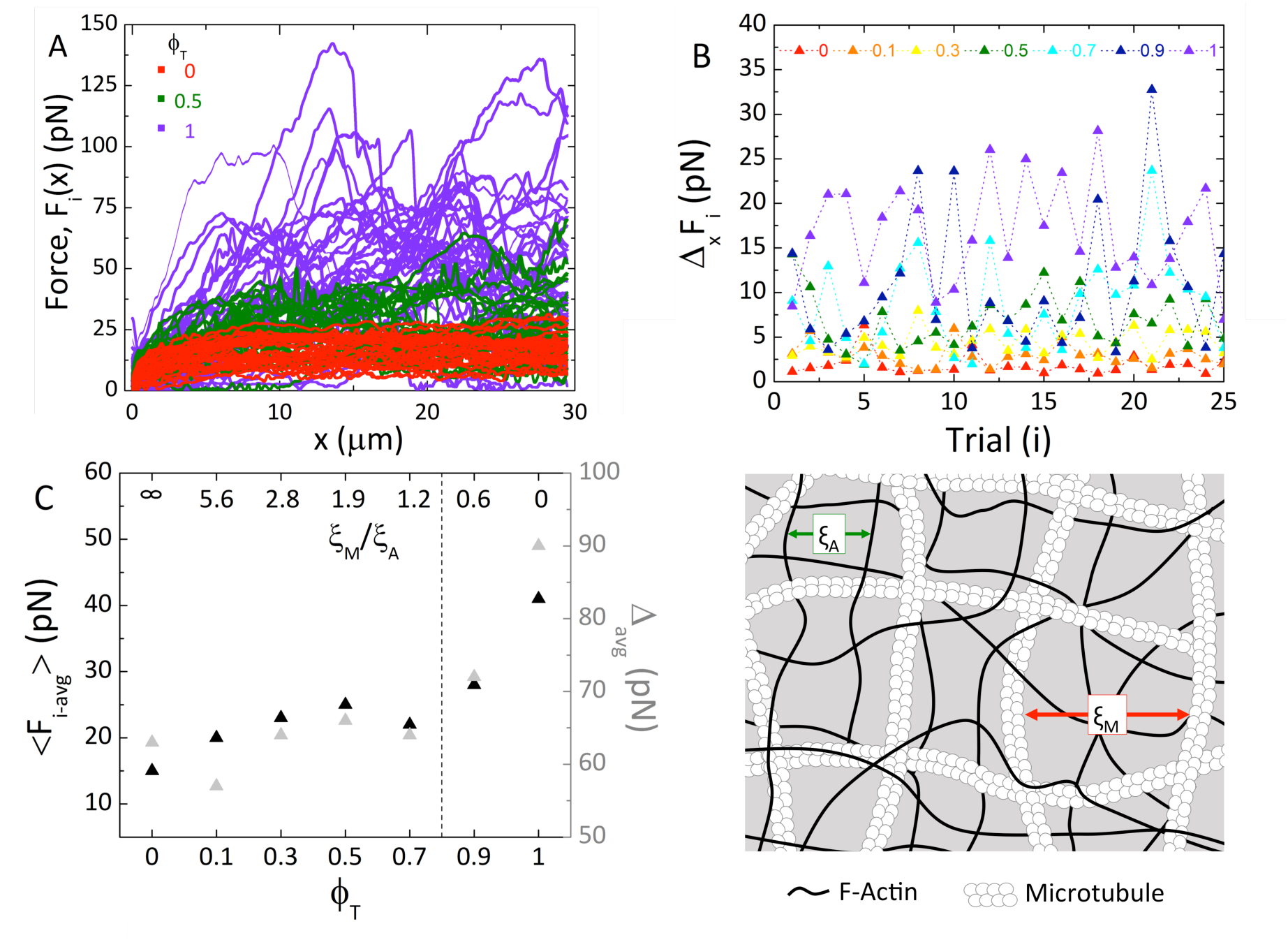
Microtubules increase average resistive force and heterogeneity of force response during strain. (A) Force curves for all 25 measurements, *i*, of composites with tubulin molar fractions of *ϕ*_*T*_ = 0 (red), 0.5 (green) and 1 (purple). Distribution of force values during strain (i.e. noise in force curves) as well as trial-to-trial variability are much higher for 100% microtubules compared to composites. (B) Standard deviation of the force values for each trial, Δ_*x*_*F*_*i*_, shows increasing microscale heterogeneity in the force response during strain with more microtubules. (C) Average force during strain, *<F_i-avg_>*, for each composite (black), as well as the corresponding percent range in strain-averaged force, Δ*_avg_ = 100*(F_i-avg___max_ – F_i -avg_min_)/(2<F_i-avg_>)* (grey) as a function of *ϕ*_*T*_ (bottom axis) and ratio of microtubule mesh size to actin mesh size *ξ*_*M*_/*ξ*_*a*_ (top axis). As shown, both the average and range in force values are surprisingly unaffected by the presence of microtubules until a tubulin fraction of 0.9. At these molar fractions (*ϕ*_*T*_ < 0.7), the mesh size of actin remains smaller than microtubules (*ξ*_*M*_/*ξ*_*a*_ *>* 1). The dotted vertical line shows when the mesh sizes for both filaments are equal (*ξ*_*m*_=*ξ*_*a*_). (D) Illustration of equimolar actin-microtubule composite (*ϕ*_*T*_ = 0.5). As depicted, the actin mesh size is *~2x* smaller than the microtubule mesh, with *ξ*_*a*_ = 0.6 μm and *ξ*_*m*_ = 1.1 μm.

To quantify the microscale heterogeneity and its dependence on tubulin fraction, we calculate the standard deviation of *F*_*i*_*(x)* values during the strain for each *i*^th^ trial, which we denote as Δ*_x_F_i_.* As shown in Fig 3B, Δ_*x*_*F*_*i*_ remains relatively small and unchanged until a tubulin fraction of *ϕ_T_ ≈* 0.7 after which the standard deviation values significantly increase. The large peaks and dips in force curves responsible for the increased Δ_*x*_*F*_*i*_ in microtubule-dominated composites are suggestive of microtubule buckling events. Previous studies have shown that microtubules buckle under large compressive loads (22, 23), except when they are integrated within the cytoskeletal matrix (20, 24, 29, 38). In line with these results, we interpret the displayed reduced standard deviation and elimination of large dips in the force during strain for *ϕ*_*T*_ < 0.7 composites as due to the elastic actin network providing reinforcement to microtubules against buckling.

To evaluate the large-scale heterogeneities, we average each *F_i_(x)* over the strain *x*, which we signify as *F*_*i-avg*_, and calculate the ensemble-averaged *F_i-avg_, <F_i-avg_>*, as well as the corresponding percent range, *A_avg_ = 100*(F_i-avg_max_ – F_i-avg_min_)/(2<F_i-avg_>).* As shown in Figure 3C we find that the average force, (*<F_i-avg_>*) is ~3*x* larger for microtubule networks than that for actin networks and the corresponding percent range (Δ_*avg*_) is doubled. However, surprisingly, the resistive force and heterogeneities do not increase smoothly as *ϕ*_*T*_ increases. Rather, they remain relatively unchanged until the tubulin fraction exceeds 0.7. To understand these results, we calculate the predicted mesh size for single-species actin and microtubule networks, *ξ*_*A*_ and *ξ*_*M*_, at each concentration present in composites. As displayed in the top axis of Figure 3C, *ξ*_*A*_/ *ξ*_*M*_ remains larger than 1 up to *ϕ*_*T*_ ≈ 0.8, with the actin mesh size ranging from ~6 to 1.25*x* smaller than that for microtubules. Only at tubulin molar fractions of 0.9 and 1 do we see a marked increase in average force and range in force as the tubulin mesh size becomes smaller than that for actin (*ξ*_*A*_/ *ξ*_*M*_ < 1). Thus, up to *ϕ_T_ ≈* 0.8 a tighter, more entangled actin network pervades the system and suppresses the heterogeneities caused by larger, more rigid microtubules. In other words, the effective mesh formed in the co-entangled composite is dominated by actin rather than microtubules until tubulin comprises ~80% of the proteins present in the composite (Fig 3C,D). Further, the “effective” composite mesh size (i.e. the smaller of the two mesh sizes, *ξ*_*A*_ and *ξ_M_)* increases as *ϕ*_*T*_ increases from 0 to 0.7. So while semiflexible actin filaments are being replaced by rigid microtubules, the decrease in filament density/entanglements offsets the increase in filament rigidity such that the resistive force remains relatively unchanged until *ϕ*_*T*_>0.7. This effect is also seen in our steady-state image analysis described below (Fig 5).

We also characterize the relaxation of induced force following strain for all composites. As shown in Figure 4A, the force relaxations for all composites display two phases of power-law relaxation in time (<*F_i_> ~ t^−α^*), an initial fast decay followed by a slow decay. Scaling exponents for the fast decay decrease roughly linearly, from *α_1_ ≈* 1.7 to 0.5, with increasing fractions of tubulin, and essentially disappear for microtubule networks (Fig 4B,C). Conversely, the exponents for the slow decay are largely insensitive to composite composition, displaying an average exponent of *α_2_ ≈* 0.4 independent of tubulin fraction (Fig 4C).

**Figure 4.**
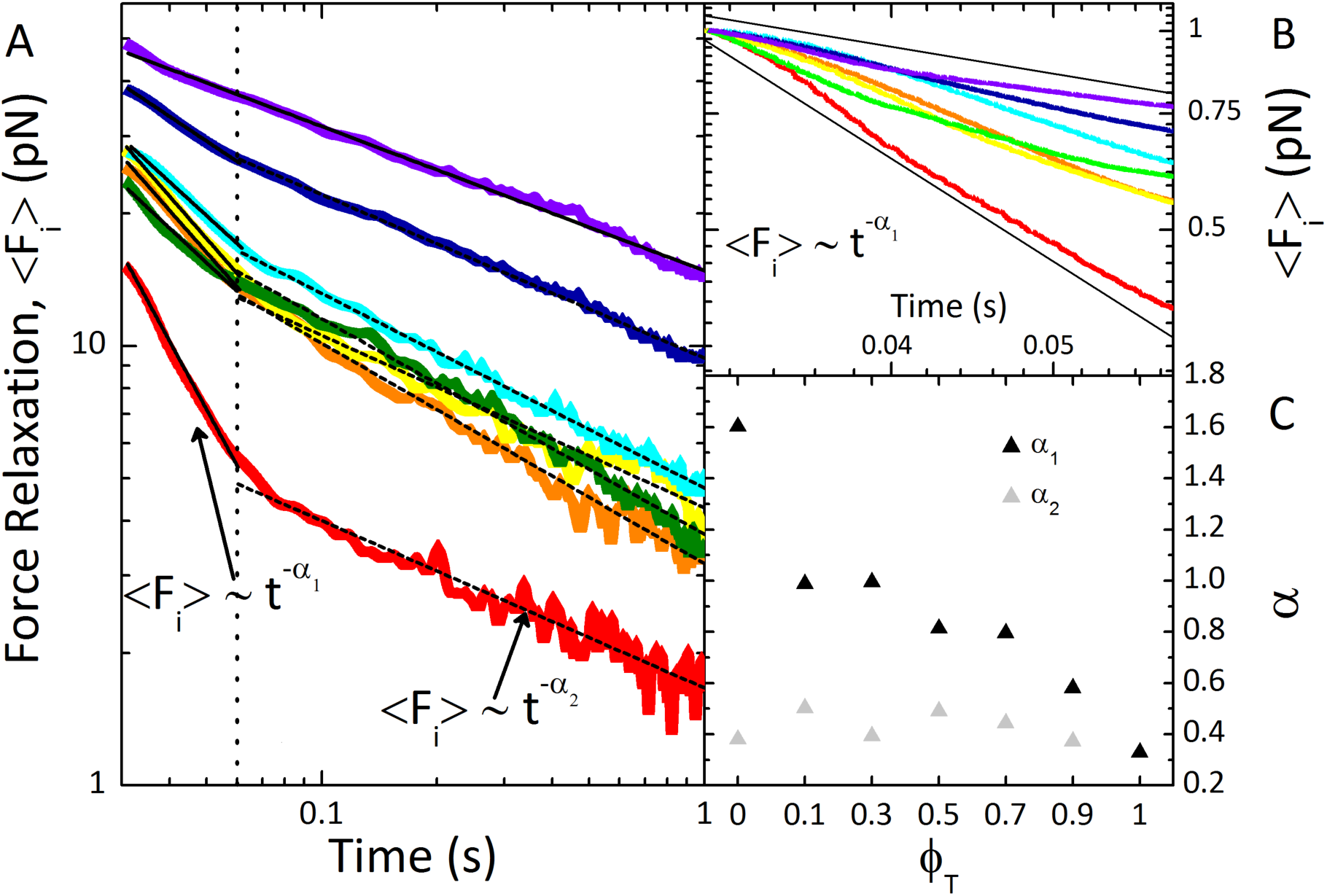
Force relaxation of actin-microtubule composites following strain exhibits two-phase power law decay. (A) Relaxation of ensemble-averaged induced force following strain as a function of time for composites of varying tubulin fractions *ϕ*_*T*_. Color scheme is as in Figures 3 and 4. Induced force relaxes via two distinct power law decays: an initial fast decay, <*F_i_> ~ t^−α_1_^*, followed by a slow decay, <*F_i_> ~ t^−α_2_^*. The vertical dotted line guides the eye to separate the two phases. The dashed lines are fits of the data to power laws with exponents *α*_*1*_ (fast decay) and *α*_*2*_ (slow decay). (B) Zoomed-in fast decay curves with black lines corresponding to scaling exponents of 1.7 and 0.4. (C) Scaling exponents measured for fast (*α*_*1*_) and slow (*α*_2_) decays. The fast scaling exponent decreases proportionally with decreasing actin concentration (i.e. increasing *ϕ*_*T*_), and becomes indistinguishable from *α*_*2*_ when *ϕ*_*T*_ = 1. Conversely, slow relaxation is independent of composite composition, with an average exponent of *α*_*2*_ ≈ 0.4.

These unique relaxation characteristics are in opposition to expected dynamics for entangled polymer systems, in which force relaxation is typically described by a sum of exponential decays due to distinct relaxation mechanisms with well-separated timescales (12, 39). The slowest of these relaxation mechanisms is predicted to be that of the entangled polymers diffusing out of deformed entanglement tubes (i.e. disengagement). However, previous studies have shown that entangled biopolymer networks can exhibit power-law force relaxation when subject to nonlinear strains (40, 41). In these studies, force relaxation transitioned from exponential to power-law when the strain rate exceeded a critical value (~2*x* slower than the rate used here). The power-law exponent of ~0.5 could be explained as arising from non-classical disengagement of entangled polymers from strain-dilated entanglement tubes (40, 42, 43). Namely, the nonlinear strain dilates the entanglement tubes, so, following strain, tubes shrink back to their original size over the timescale that filaments reptate out of the deformed tubes. The result is that, as each filament attempts to reptate out of its tube, the characteristic disengagement time grows longer, thereby producing power-law decay. Our second relaxation phase, with scaling exponent close to the previous finding of 0.5 for actin networks, can likewise be explained by this non-classical disengagement mechanism (40). Further, the insensitivity of *α*_*2*_ to *ϕ*_*T*_ indicates that entanglements rather than filament flexibility drive the relaxation.

Unlike the slow relaxation rate, the fast relaxation has a clear dependence on composite composition. As *α*_*1*_ scales roughly linearly with the fraction of actin in composites, this fast mode likely arises from actin filaments in the composite. In contrast to rigid microtubules, entangled actin can partially relax induced stress via bending fluctuations, on faster timescales than that of disengagement (12, 14). However, introducing microtubules can reduce these non-affine bending deformations (15), which can also promote stress-stiffening (as seen in Fig 2). Because composites exhibit a clear transition to stiffening behavior as *ϕ*_*T*_ increases, we can understand this fast relaxation mode as arising from actin bending fluctuations that microtubules increasingly suppress as *ϕ*_*T*_ increases.

To correlate the measured nonlinear stress characteristics with the corresponding steady-state dynamics we quantify the mobility of actin and microtubules in steady-state composites by analyzing time-series acquired using two-color fluorescence confocal microscopy (Fig 5). As described in the Materials and Methods, we use the standard deviation of pixel intensities over time (SD) as a measure of mobility. As shown in Figure 5, the mobility of both actin and microtubules within composites display a surprising non-monotonic dependence on tubulin fraction: mobility increases as *ϕ*_*T*_ increases to 0.5 after which mobility is reduced. The drop in mobility occurs at the same tubulin fraction at which composites transition from stress-softening to stiffening, further demonstrating that stiffening indeed arises from microtubules suppressing filament fluctuations. The increase in mobility as *ϕ*_*T*_ increases to 0.5 is less intuitive as semiflexible actin filaments are being replaced with more rigid microtubules. However, while the composite is becoming more rigid the “effective” mesh size actually increases with *ϕ*_*T*_ until *ϕ*_*T*_ ≈ 0.7, allowing both actin and microtubules to more easily move. This increasing mobility also aligns with our finding that the force response is largely insensitive to *ϕ*_*T*_ until ~0.7 (Fig 3C). While network rigidity increases, which would increase resistive force, the mesh size is also increasing, allowing filaments to more easily move to alleviate stress, offsetting this resistance.

**Figure 5.**
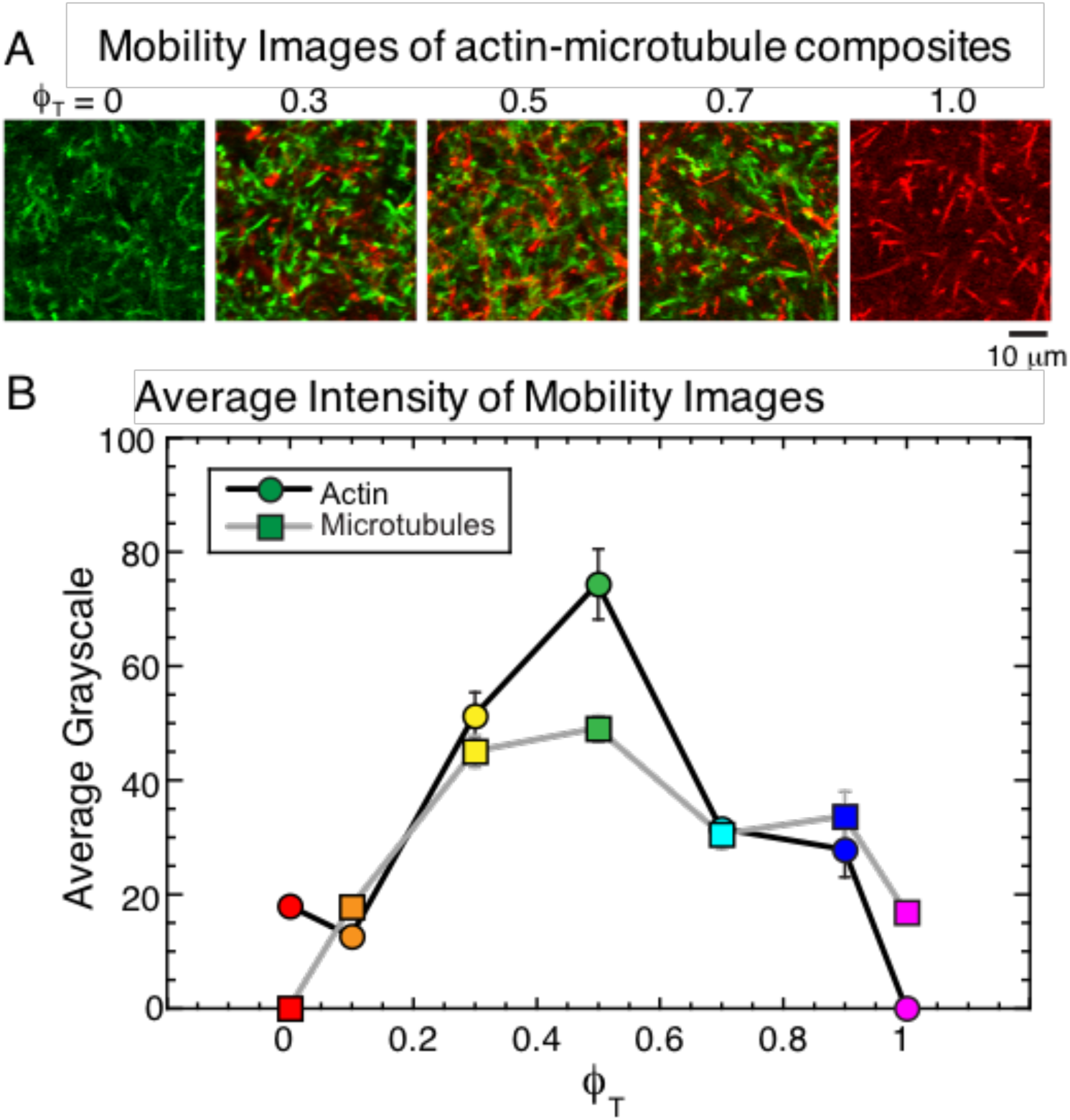
Mobility of both actin and microtubules in composites exhibit a non-monotonic dependence on tubulin fraction. (A) Each image displays the temporal standard deviation of intensity values for each pixel of a two-color confocal time-series (180 frames, 1 fps). Images show the standard deviation for actin (green) and microtubules (red) in a 44 µm^2^ area for composites of varying tubulin fractions (*ϕ*_*T*_) listed below the corresponding image. As described in Materials and Methods, these “mobility images” show the extent to which filaments move, with higher pixel intensities signifying more motion. (B) The average of all standard deviation intensity values from mobility images vs tubulin fraction *ϕ*_*T*_. Error bars are the standard error from averaging over 512×512 pixels in 3–5 images for each *ϕ*_*T*_. As shown, the mobility of both actin (circles) and microtubules (squares) increases as *ϕ*_*T*_ increases to 0.5, followed by a subsequent drop.

## Conclusion

We use optical tweezers microrheology and two-color fluorescence microscopy to characterize the mesoscale force response and mobility of co-entangled composites of actin and microtubules, which we create by *in situ* co-polymerization of actin monomers and tubulin dimers using custom-designed hybrid conditions. By systematically varying the relative concentrations of actin and microtubules (quantified by the molar fraction of tubulin *ϕ*_*T*_), we show that composites exhibit a wide range of dynamical properties that can be tuned by *ϕ*_*T*_. Our collective results demonstrate that microtubules suppress actin bending fluctuations, enabling composites to stiffen in response to strain and relax induced force more slowly; while actin supports microtubules against buckling by providing a soft semiflexible mesh that permeates the larger microtubule mesh and partially absorbs induced stress. We also show that the interplay between varying mesh sizes and filament rigidity leads to emergent fast mobility in equimolar composites compared to networks of mostly actin or microtubules. These results demonstrate the synergistic ways in which thin, semiflexible actin filaments and thick, rigid microtubules can sterically interact to enable the myriad of mechanical processes and properties the cytoskeleton exhibits. Our quantitative analysis of the dependence of mechanical properties and mobility on composite composition also provides important new insights into how composite materials can be tuned to display user-defined mechanics, and how cells can likewise tune their mechanical properties by using varying cytoskeletal filament networks. This study provides the groundwork for future investigations on in vitro cytoskeletal composites that include crosslinking proteins, intermediate filaments, and motor proteins. These future studies will expand the phase space of mechanical properties that these bio-inspired composite materials can exhibit, and shed light onto the role that each of these additional components play in the mechanics of the cytoskeleton.

## Author Contributions

S.N.R. designed experiments, conducted experiments, analyzed data and wrote the manuscript; J.L.R. designed experiments, analyzed and interpreted data, and helped write the manuscript. R.M.R.-A. designed experiments, analyzed and interpreted data, and wrote the manuscript

## Acknowledgements

This research was funded by a Scialog Collaborative Innovation Award from Research Corporation and the Gordon & Betty Moore Foundation (grant no. 24192, awarded to RMR-A and JLR); an NSF CAREER Award (grant no. 1255446, awarded to RMR-A);

